# Language modelling for biological sequences – curated datasets and baselines

**DOI:** 10.1101/2020.03.09.983585

**Authors:** Jose Juan Almagro Armenteros, Alexander Rosenberg Johansen, Ole Winther, Henrik Nielsen

## Abstract

**Motivation:** Language modelling (LM) on biological sequences is an emergent topic in the field of bioinformatics. Current research has shown that language modelling of proteins can create context-dependent representations that can be applied to improve performance on different protein prediction tasks. However, little effort has been directed towards analyzing the properties of the datasets used to train language models. Additionally, only the performance of cherry-picked downstream tasks are used to assess the capacity of LMs.

**Results:** We analyze the entire UniProt database and investigate the different properties that can bias or hinder the performance of LMs such as homology, domain of origin, quality of the data, and completeness of the sequence. We evaluate n-gram and Recurrent Neural Network (RNN) LMs to assess the impact of these properties on performance. To our knowledge, this is the first protein dataset with an emphasis on language modelling. Our inclusion of properties specific to proteins gives a detailed analysis of how well natural language processing methods work on biological sequences. We find that organism domain and quality of data have an impact on the performance, while the completeness of the proteins has little influence. The RNN based LM can learn to model Bacteria, Eukarya, and Archaea; but struggles with Viruses. By using the LM we can also generate novel proteins that are shown to be similar to real proteins.

**Availability and implementation:** https://github.com/alrojo/UniLanguage

## 1 Introduction

The task of language modelling (LM) (Bengio *et al.*, 2003; Merity *et al.*, 2017) has proven a successful strategy to build unsupervised contextual representations of text leading to state-of-the-art performance in many supervised natural language processing (NLP) tasks (Devlin *et al.*, 2018; Peters *et al.*, 2018).

Protein sequences are in many ways comparable to text: discrete symbols (amino acids), dictionary of up to 25 symbols (similar to characters), average length of 335 (like a paragraph), and access to large databases of unlabelled sequences (akin to English Wikipedia).

Recent work has shown that LMs trained on protein sequences can be used to improve performance on multiple protein prediction tasks (Heinzinger *et al.*, 2019; Rives *et al.*, 2019). However, in contrast to NLP where the field of language modelling has been studied in 20+ years before being used for pretraining, little effort has been directed towards studying the desired properties of datasets and methods for language modelling on proteins.

As a result, current methods naively train LMs without considering the vast heterogeneity of proteins in nature, evidence for protein existence, protein fragments, and homology partitioning of training, validation, and test sets. Mixing the taxonomic domains of life can be compared to training a single LM on several different languages at the same time, potentially missing out on domain specific specialization. This can make it particularly challenging to learn underrepresented domains. Protein fragments might be incomplete, discontinuous, or filled with unknown symbols. Lack of experimental evidence, which constitutes 99% of UniProt, can cause noisy training and unreliable testing. Not considering homology partitioning can cause proteins with high similarity to be placed in different partitions resulting in an overestimation of generalization performance potentially leading to erroneous conclusions. Moreover, many homologous sequences within a training set might lead training to overly focus on a certain family of proteins or misjudge the true amount of unique proteins.

Figure 1 highlights the representation of taxonomic domains in UniProt, the distribution of proteins across dataset qualities, the amount of protein fragments, and homology overlap in the experimental dataset. It can be observed that most of the data is predicted (no experimental evidence of existence), Bacteria is overly represented, and Viruses have a low amount of unique proteins.

**Fig. 1:**
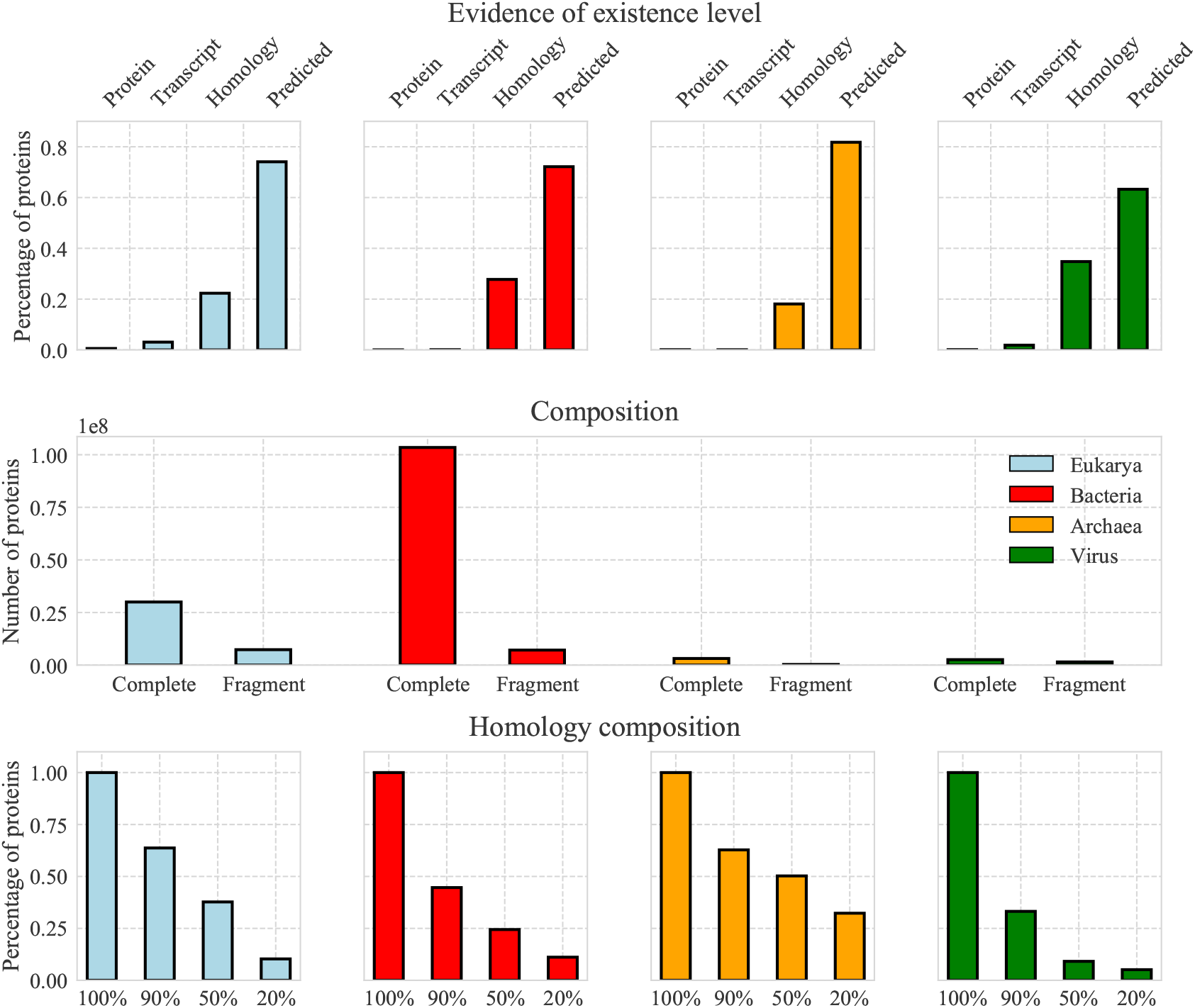
UniProt composition based on quality, completeness and homology. The different properties are divided by domain, where each color correspond to a different domain. The evidence of existence is divided in four levels, where Protein and Transcript (high quality) have experimental evidence while Homology and Predicted lack experimental evidence (low quality). The composition of the data is divided in complete sequences and fragments, which is described in hundreds of millions. The homology composition represents the proportion of experimental proteins that remain when the dataset is homology reduced by four different similarity thresholds. This indicates the degree of redundancy in the dataset.

We contribute a new language modelling dataset for protein sequences. This dataset addresses all the above-mentioned concerns and provides a language modelling dataset with homology partitioning; only high quality samples for validation and testing; and segmentation according to domain, quality, and completeness.

The end-goal with language modeling is to develop contextual representations encapsulating important protein features. With such contextual representations, we believe that state-of-the-art for all known protein prediction tasks can be improved. Moreover, with understanding of protein likelihood we can filter, auto-complete and generate new novel proteins. However, this end-goal requires that language models are capable of producing great contextual understanding of protein sequences from all domains of interest. Measuring perplexity is commonly used in natural language processing for evaluating contextual representations. Perplexity is a measure of how well a language model can predict a protein. Perplexity allows us to evaluate context-dependent representations without the need of supervised protein tasks. This makes perplexity easily applicable and allows us to evaluate across the entirety of UniProt, which is why we use it for protein evaluation.

Our hope with this dataset is to facilitate language modeling and understand how the varying levels of evidence for proteins (quality), different taxonomic domains, and protein completeness impact the ability to find regularity in protein sequences.

We evaluate a baseline *n*-gram and a modern recurrent neural network architecture known as the AWD-LSTM. We train the baseline and AWD-LSTM on varying levels of data quality (high quality, low quality, both), the inclusion of taxonomic domains (training on one domain vs. all domains), and only using incomplete proteins. When evaluated all models on the high quality validation and test sets for all domains, we find that that domain and dataset quality have a significant effect on perplexity. Moreover, when optimizing the AWD-LSTM hyperparameters we found that default NLP hyperparameters often underperformed, achieving equivalent perplexity to a count-based unigram model. To find the optimal parameters, we had to do an extensive hyperparameter search using bayesian hyperparameter optimization tools (Hayes *et al.*, 2019).

In conclusion, we have defined a massive, curated, and standardized dataset for language modelling on protein sequences. We provide results from *n*-gram and modern recurrent neural network models.

## 2 Related work

In recent years significant progress has been made in language modelling of text (Bengio *et al.*, 2003). Most noticeable is the distributed encodings of hidden states using deep neural networks, which can learn to compress, understand, and produce sentences in fluent English (Mikolov *et al.*, 2010; Merity *et al.*, 2017). Current state-of-the-art within language modelling is based on attention architectures (Bahdanau *et al.*, 2014; Vaswani *et al.*, 2017; Dai *et al.*, 2019) and the access to immense computing resources and large datasets (Radford *et al.*, 2018, 2019). It has been found that these large language models have a profound impact on generating contextual embeddings for NLP tasks (Peters *et al.*, 2018; Radford *et al.*, 2018).

Language modelling is unidirectional as it decomposes the joint probability. However, text, like proteins, is often given in a complete form, presenting a possible bidirectionality that cannot be captured by unidirectional language models. As a response, the models BERT (Devlin *et al.*, 2018) and XLNet (Yang *et al.*, 2019) provide bidirectional language modeling objectives by taking inspiration from denoising autoencoders (Vincent *et al.*, 2008), which currently rank state-of-the-art on popular benchmarks such as GLUE (Wang *et al.*, 2018).

However, these state-of-the-art models are prohibitively expensive to train. Because of such, we use the computationally less demanding Average Stochastic Gradient Descent Weight-Dropped LSTM (AWD-LSTM) (Merity *et al.*, 2017) for our benchmarks, which has comparable results to state-of-the-art in unidirectional language modeling. For future work we encourage researchers to test out modern attention architectures and bidirectional language modelling objectives.

In the literature for language modeling on proteins, Rives *et al.* (2019) performed language modelling on all of UniProt and did not consider homology overlap. Heinzinger *et al.* (2019) and Alley *et al.* (2019) improved upon this by using UniRef50, which is UniProt with 50% homology reduction. However, 50% identity still implies a very significant homology between proteins (Sander and Schneider, 1991). They randomly assigned dataset partitions, which resulted in most of their test set not being experimentally validated. Moreover, none of the mentioned papers considered the domain of origin or protein completeness. The difference between taxonomic domains, and bias towards highly represented domains such as Bacteria, can cause the models to be biased.

When evaluating language model performance, most recent work has copied the style from natural language processing by reporting scores on supervised tasks using their contextual representations. Rao *et al.* (2019) propose a combination of five protein prediction tasks to assess the performance of contextual representations for protein sequences. However, the performance of supervised tasks is not a direct measure of how informative contextual representations are and suffers from dataset biases. Perplexity on the other hand directly measures the models ability to predict amino acids, which reflects the model capability to create informative contextual representations. Nonetheless, Rao *et al.* (2019) can provide an ability to assess if the contextual representation captures certain structural phenomena, which perplexities will not provide.

## 3 Methods

Language modeling is the task of estimating the joint probability distribution over a finite sequence of length *l* of discrete tokens:

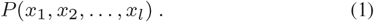

A token, 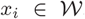, comes from a size *k* discrete dictionary, 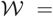 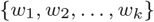.

By modelling the probability of a sequence we can evaluate whether a sentence is likely to exist. Defining the dictionary, 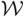, as the set of amino acids allows us to calculate the probability of a protein sequence existing.

As equation (1) is hard to calculate we will introduce two methods, popularized in NLP, for doing language modeling of proteins: The *n*-gram language model and the Recurrent Neural Network Language Model (RNN-LM). To test language modeling capability we propose the biggest known language modelling dataset, with homology partitioning, by clustering the experimental part of UniProt. The dataset consists of different subsets in order to test the influence on language modelling of taxonomic domain, quality, homology, and protein completeness.

### 3.1 Conditional language modelling

The *n*-gram and RNN-LM models both rely on rewriting eq. (1) using the chain rule of probability:

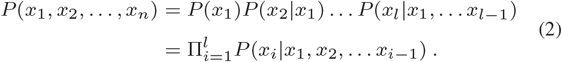

In order to make a model that scales, the *n*-gram makes a Markov assumption and the RNN-LM uses a learnable compression of the conditionals.

### 3.2 *n*-gram models and the Markov assumption

The *n*th order Markov assumption relaxes the rules of conditional probability by assuming that predicting the next token only depends on the *n* preceding tokens:

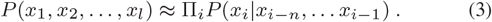

This is a strong assumption that is definitely not true for neither text nor proteins. However, it can be argued that some tokens have a larger influence on predicting future tokens. In particular for text, the most recent words are highly influential on what might come next.

Using the chain rule we can write:

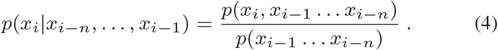

In a pure maximum likelihood, frequencies are used to estimate probabilities so the training set counts can be used to estimate the conditionals:

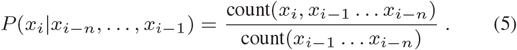

The simple *n*-gram technique has been shown to work surprisingly well in NLP literature when calculating the probability of text. Laplace smoothing, also known as the add-one trick, where count → count +1 is used as regularization to better deal with low count *n*-grams (Jurafsky and Martin, 2009).

### 3.3 Recurrent Neural Network Language Models

The Recurrent Neural Network Language Models (RNN-LM) is a neural network based language model that circumvents the statistical based approach of the n-gram model by compressing the conditional into a hidden state, *h*_*i*_ ∈ ℝ^*d*^, using neural networks:

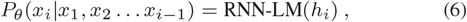

where the RNN-LM is a non-linear, potentially multi-layered, function that outputs the probability of *x*_*i*_. The RNN-LM repeatedly updates the the hidden state, *h*_*t*_, at every time step, allowing the RNN-LM to predict the probability of a sequence in linear time w.r.t. the sequence length

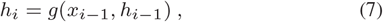

where *g* corresponds to the recurrent neural network portion of the RNN-LM. We use the AWD-LSTM (averaged stochastic gradient descent weight dropped LSTM) as our specific RNN-LM implementation (Merity *et al.*, 2017). The AWD-LSTM has several hyperparameters that we optimize with SigOpt (Hayes *et al.*, 2019).

### 3.4 Inference

Given a trained RNN-LM model, we can generate novel protein sequences by iteratively sampling from the network’s output distribution, 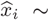 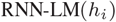, then feeding in the sample as input at the next timestep 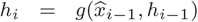. While the RNN-LM is deterministic, the stochasticity of multinomial sampling gives a distribution over protein sequences. As sampling is conditional on the hidden state, the distribution of sequences depends on previous inputs. This allows the RNN-LM to complete any partial sequence. For generating a completely novel sequence, we will provide the RNN-LM with a hidden state, *h*_1_, of zeroes and *x*_1_ set to Methionine.

### 3.5 Evaluation metric

Historically, perplexity has been the preferred method to evaluate model performance in language modelling literature (Jurafsky and Martin, 2009; Dai *et al.*, 2019; Merity *et al.*, 2017). Perplexity is the exponential of the average log-likelihood and measures how well a language model predicts a sequence of amino acids.

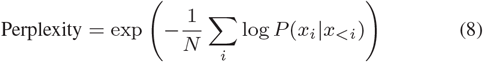

Because of the log term very confident and wrong predictions will weight more. When reporting perplexities we will consider the protein sequences as one contiguous sequence, such that long proteins will have a larger influence. Notice, that predicting uniform probability over all classes will correspond to a perplexity equal the amount of classes. In our case, the uniform perplexity will be 26 for our 25 amino acids + end-of-sequence symbol.

### 3.6 Datasets

The data used in this project are downloaded in their entirety from the UniProt database (UniProt Consortium, 2019). For each protein, we extract information regarding its experimental evidence (experimentally validated or predicted), domain of origin (Eukarya, Bacteria, Archaea, or Viruses), and completeness (i.e. whether the protein is complete or fragmented). This gives in total 2 × 4 × 2 = 16 datasets. The exact definition of these criteria are discussed below and summarized in Table 1.

**Table 1.** Number of sequences and distribution of lengths for each domain and protein quality.

|  |  | Eukarya | Bacteria | Archaea | Viruses |
| --- | --- | --- | --- | --- | --- |
| Experimental | Mean length | 428 | 489 | 327 | 376 |
|  | High quality | 899K | 69K | 3.3K | 30K |
|  | Low quality | 421K | 7916 | 2252 | 30K |
|  | Proteins removed | 52K | 43K | 67 | 22K |
| Predicted | Mean length | 460 | 313 | 280 | 365 |
|  | High quality | 17.9M | 56.9M | 1.9M | 1.0M |
|  | Low quality | 4.3M | 3.6M | 188K | 1.1M |
|  | Proteins removed | 17.6M | 68.5M | 1.8M | 678K |

The dataset is divided into two sets depending on the evidence of their existence using the field *Protein existence* in UniProt. Proteins can have *experimental* evidence at protein and transcript level. Proteins with existence being inferred by homology or predicted by gene finding methods are regarded as *predicted* proteins.

Fragmented proteins are incomplete protein sequences that might be from a random place in the original sequence, contain many unknowns, or noncontiguous regions. We also consider proteins that do not start with Methionine as fragments.

The high quality dataset is homology partitioned into training, validation, and test set. We employ 60% for training, 10% for validation and 30% for testing. When partitioning we first cluster the proteins based on a similarity threshold of 20% identity. This means that proteins with an identity higher than this threshold are grouped in the same cluster. For the clustering task, we utilize the MMSeqs2 tool (Steinegger and Söding, 2017). Each cluster is then assigned to one of the three possible partitions, taking into consideration that the proportions of taxonomical domains, and fragmented proteins, are the same across all partitions.

The low quality dataset is aligned to the validation and test sets obtained from the high quality homology partitioning. MMSeqs2 is also used for this alignment process. Low quality proteins with a similarity above 20% identity are discarded. The remaining proteins are kept as the low quality training set. For both high and low quality datasets, we remove all duplicated proteins. The dataset statistics can be found in Table 1.

## 4 Experimental Design

In our experiments we test the AWD-LSTM (Merity *et al.*, 2017) and *n*-gram model with add-one smoothing (Jurafsky and Martin, 2009) on the datasets defined above in Section 3.6.

### 4.1 Model details

As our specific RNN-LM implementation, we modify the AWD-LSTM by using SGD instead of averaged SGD. We reduce the learning rate if there has been no improvements over 10 epochs for the experimental training sets. If the learning rate was reduced more than 5 times, the training was stopped. For the predicted datasets we use iterations comparable to 10 epochs on the experimental eukaryotes (about 600k samples per epoch). To optimize the hyperparameters of the AWD-LSTM we use SigOpt (Hayes *et al.*, 2019) and define the parameter space as follows: learning rate [1, 20], learning rate decay size [0.05, 1], batch size [32, 256], back propagation through time [32, 256], LSTM layers [2, 5], number of hidden LSTM units in each layer [128, 1280], gradient clipping [0.05, 1], weight decay [0.0, 0.001], all dropouts individually set [0, 0.6].

On the experimental dataset we train and validate 300 models on Eukarya. We then take the best performing hyperparameters of the Eukarya and use it on all other language modelling training sets to achieve comparable results. We also tried to optimize AWD-LSTM parameters individually for each domain on the experimental training set. Here we found that only Archaea benefits from having individually optimized parameters, which we comment on in the result section.

Each model is trained on a single GTX-Titan-X GPU with 12 Gb of internal memory. We use the Salesforce implementation of AWD-LSTM^1^, which is written in the PyTorch framework (Paszke *et al.*, 2017).

### 4.2 Quantitative evaluation

Given a homology partitioned validation and test set of experimental quality and identical model hyperparameters, we compare the perplexities of different datasets. We train on single domain experimental, predicted, and combined data; on all-domain experimental and combined data; and finally on fragments for Eukaryotes.

### 4.3 Qualitative evaluation

To better understand what the model learns and when it is confident in its predictions, we analyse model likelihood and proteins that have been generated by the AWD-LSTM. We take a single protein and overlay the amino acid probability with regions that have known secondary structures (non-coil). To assess whether proteins generated by the trained model has protein-like features, we use SignalP 5.0 (Almagro Armenteros *et al.*, 2019) to predict signal peptides in proteins generated by the AWD-LSTM model. We compare their signal peptide logos with logos generated from real proteins and from proteins generated by a baseline model.

## 5 Results and Discussion

We present quantitative results with language model performance on the experimental test set of each domain. For comparison, we evaluate *n*-gram models, and present results from training the AWD-LSTM on varying dataset qualities, domains, and completeness.

### 5.1 *n*-grams baseline

The *n*-gram models are used as a baseline in language modeling in protein sequences, results are shown in Figure 2. This performance varies for the different taxonomic domains, with Viruses being the worst performing domain while Archaea achieve the lowest perplexity. Regarding the size of the *n*-gram context, a bigram and trigram achieve the best performance, with the exception of Viruses, where a unigram (predicting according to amino acid frequencies) obtains the best performance. The best perplexity for the different domains are: Eukarya 18.09, Bacteria 17.49, Archaea 17.31, and Viruses 18.83. Anything below the perplexity of the unigram indicates that the model has found structure in the sequence of amino acids.

**Fig. 2:**
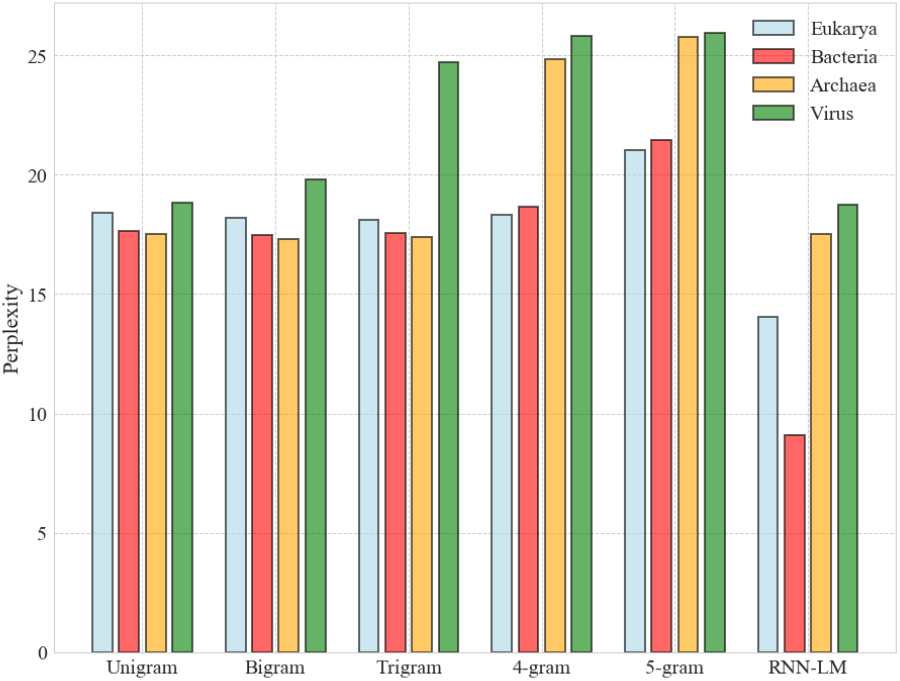
*n*-gram and RNN-LM perplexity test performance (low is better) for different *n* and domains of life. The *n*-gram is calculated on the experimental training set separately for each domain.

### 5.2 Quality and origin of training data

The RNN-LM performance can be found in Table 2. We use the same model for all dataset combinations. The model is described in Section 4.1. The experimental column can be directly compared with the *n*-grams as the training and test setup is identical. We can observe that for Eukarya and Bacteria, the perplexity is significantly lower, with a reduction of 3.87 and 8.22 perplexity, respectively. This indicates that the RNN-LM is able to learn the contextual information preceding each amino acid. However, the perplexity of the RNN-LM for Archaea and Viruses are equal to the unigram perplexity. This indicates that the RNN-LM has been unable to capture sequential information in the amino acid sequence. The most reasonable explanation for these results is the minuscule amount of data used to train Archaea and Viruses. Together with the high complexity of the RNN-LM, it can lead to a quick overfitting on the training data. Nonetheless, when optimizing the hyperparameters for each domain separately, we surprisingly found that the best hyperparameters for Archaea gave a perplexity of 10.5, while optimizing for virus gave no improvement over unigram.

**Table 2.**
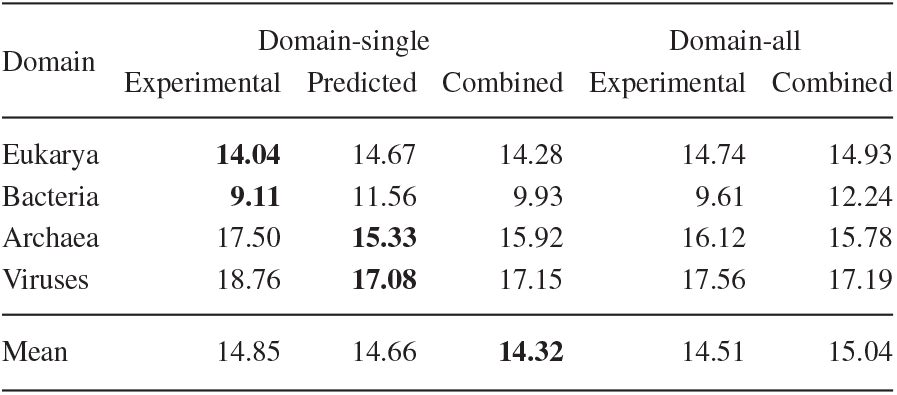
Model test performance when trained separately for each domain and for all the domains simultaneously. Test performance is in all cases measured on experimental test set. Test results for models trained with the RNN-LM. In domain-single, each row and column pair corresponds to a model trained with a specific domain and existence class and then tested on the same domain. E.g. Bacteria predicted is trained on predicted Bacteria sequences and then tested on the experimental Bacteria test set (as described in section 3). In domain-all, each column corresponds to one model trained on all domains for a specific dataset quality and then tested for each domain individually.

The lack of experimental data in Archaea and Virus is more apparent when looking at the performance of RNN-LM trained on the predicted protein data. The predicted data, even though of unverified existence, has a significantly greater amount of examples. Here we can see that the perplexity for both Archaea and Viruses is lower than when training only on the experimental dataset.

For Eukaryotes and Bacteria we observe that RNN-LMs trained on predicted dataset exhibit a slightly higher perplexity than when trained on the experimental dataset. This finding is striking considering that there is approximately 20 and 900 times more data in the predicted dataset for Eukarya and Bacteria, respectively.

These results demonstrate that the evidence of existence of protein data impacts the perplexity of protein LMs. We find that predicted datasets, even though high quantity, are not as reliable as the experimental data and introduce noise in training. However, this higher perplexity could also be due to other reasons. One possibility could be that some of the organisms represented in the predicted data are phylogenetically distant from the organisms represented in our experimental test set. This can lead to a biased model that favours the amino acid structure of the most represented organisms used to train the model. Moreover, the predicted and combined results should be very similar as we do not oversample the experimental proteins. Though, we still see a modest difference in their results, this could be due to seed and optimization variance.

### 5.3 Training on all domains

In Table 2 we show the results obtained when training a model on the data from all domains. The objective of this analysis is to address the question of whether a single RNN-LM can simultaneously learn to predict proteins belonging to different domains. The performance for the four different domains is comparable to the performance of the RNN-LM trained individually on each domain. The perplexity is slightly higher in the combined RNN-LM, but not by a large margin. This suggests that the RNN-LM is powerful enough to model multiple distributions of proteins at once. Another possibility could be that proteins from different domains share some properties that are easily captured by the RNN-LM. We can conclude that as only small differences are found between individual and combined training, the choice of training would be dependent on the downstream task usage. Furthermore, if the sole objective is to determine the probability of a protein belonging to one of these domains, then a domain-specific LM would be preferred.

### 5.4 Cross-domain predictions

We investigated whether a domain-specific RNN-LM is able to distinguish between proteins from different taxonomic domains. We hypothesized that domain-specific models would assign a higher perplexity to proteins belonging to different domains. In Table 3 we present results where an RNN-LM is trained on a single domain and evaluated on all domains individually. We found that in most of the cases, a domain-specific LM assigns a higher perplexity to proteins from other domains. One clear example is the case of Eukaryotes and Bacteria. While a model trained on Eukaryotes attains a lower perplexity on bacterial proteins (12.83) than on its own domain (14.04), the bacterial perplexity is still much higher than a Bacteria-specific RNN-LM (9.11). This could also indicate that bacterial proteins are easier to predict than eukaryotic proteins. The performances obtained by Archaea and Viruses do not support this finding due to the reduced amount of training data. For this purpose, we also looked at the performance obtained by an RNN-LM trained on predicted protein data, which is shown in Table 4. In this case, we find that a domain specific model is best at predicting its own domain. Interestingly, Viruses are very hard to predict, as even the Virus-specific model has lower perplexities on other domains than itself.

**Table 3.** Extrapolation when only training on experimental data of one domain (row) and testing on other domains (column). E.g. Eukaryotes-Bacteria means that the model was trained on Eukaryotes and tested on Bacteria.

| Domain | Eukarya | Bacteria | Archaea | Viruses |
| --- | --- | --- | --- | --- |
| Eukarya | **14.04** | 12.83 | 16.48 | **17.58** |
| Bacteria | 17.74 | **9.11** | **16.30** | 18.17 |
| Archaea | 19.08 | 17.83 | 17.50 | 19.29 |
| Viruses | 18.62 | 18.06 | 18.24 | 18.76 |

**Table 4.** Cross-domain prediction when training on predicted data of one domain (row) and testing on experimental data of other domains (column).

| Domain | Eukarya | Bacteria | Archaea | Viruses |
| --- | --- | --- | --- | --- |
| Eukarya | **14.67** | 12.70 | 16.51 | 17.36 |
| Bacteria | 16.87 | **11.56** | 16.18 | 17.72 |
| Archaea | 16.71 | 15.03 | **15.33** | 17.68 |
| Viruses | 16.64 | 16.02 | 16.71 | **17.08** |

### 5.5 Incomplete or fragmented proteins

Finally, we investigate the impact of training with proteins that are either incomplete or fragmented. We only consider fragments belonging to experimental data of Eukarya as it is the most abundant set of high quality fragments. We find that the performance on the Eukarya full sequence test set is 14.66. This result is unexpected to us, as we believed that fragments of proteins would introduce undesired noise to the real distribution of amino acid sequences. However, it seems that this is not the case and a model trained on fragments can perform almost as well as a model trained on full proteins. One possible explanation could be that the information to predict the next amino acid is more local and only short-range dependencies are required. In this context, a fragment would be equivalent to a full protein, as long-range dependencies in full sequences would be underutilized.

### 5.6 Performance over sequence length

Additionally, we studied whether the length of the protein has an influence on the overall protein perplexity. We analyze the perplexity of eukaryotic proteins in relation to their sequence length. Figure 3 illustrates that short proteins are more challenging for the RNN-LM. In particular, protein sequences below 150 amino acids have a higher perplexity. This result suggests that perplexity decreases as the protein length increases, which could indicate that longer sequences are either easier to predict or that the RNN-LM can better exploit the larger context of the protein.

**Fig. 3:**
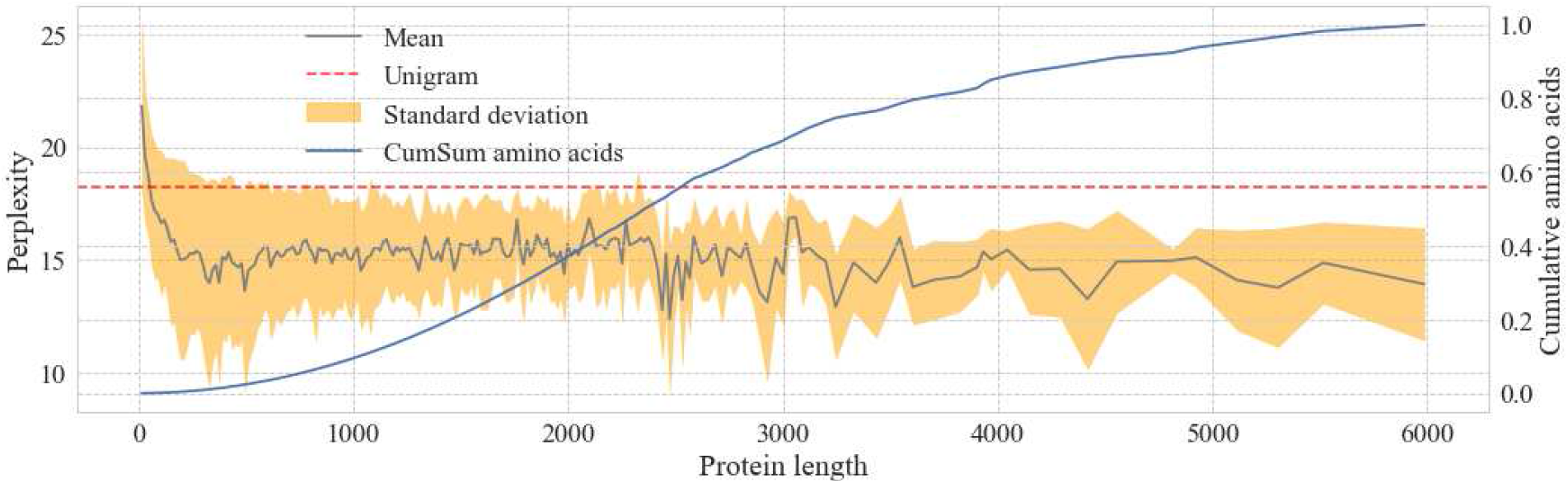
Perplexity of eukaryotic proteins by sequence length. The plot represents the mean and standard deviation of the perplexity for each protein length. The unigram perplexity is included as a baseline. The cumulative sum of the amino acids at each length interval is also represented.

### 5.7 Qualitative analysis

#### 5.7.1 Probabilities along sequence

When executing the RNN-LM over a protein sequence we obtain a likelihood for each residue. This likelihood estimate can be used to assess the model’s abilities at specific positions, or regions, in the sequence. One example of likelihood over the sequence is illustrated in Figure 4. In this example, we can observe that certain regions have a high likelihood, even close to 1. When comparing these regions against the amino acids with secondary structure we see an overlap. This suggests that the model is more correct and confident in regions with secondary structure, as these regions are generally more conserved than the rest of the sequence. For future work, it would be of interest to further analyse the correlation between the likelihood of the RNN-LM and secondary structure and/or conservation.

**Fig. 4:**
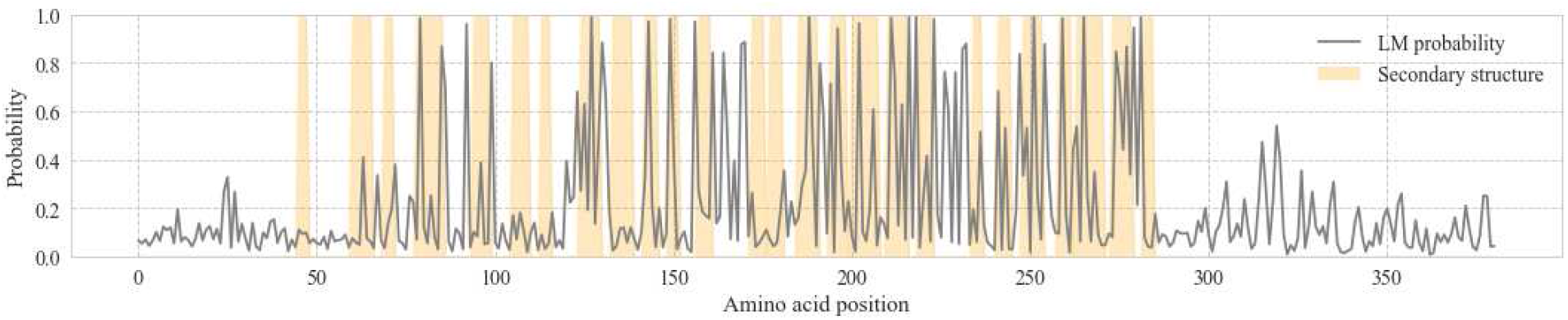
Probability per residue along the sequence of the CD55 human protein. Secondary structure elements are overlaid.

#### 5.7.2 Discriminating between real and random proteins

One of the RNN-LM outputs is the perplexity of the overall protein sequence. This perplexity can be interpreted as a score of how much the protein resembles the training set – i.e. how much it resembles a real protein. To better understand how well the RNN-LM performs we compare the average protein perplexity over our test set to a randomly generated sequence of amino acids. The random sequences are sampled with the underlying frequencies of amino acids of eukaryotic proteins (unigram probability). Figure 5 shows the perplexity assigned by the RNN-LM to experimental eukaryotic proteins and unigram sampled proteins. We can observe that, even though both sets have the same amino acid composition, the RNN-LM assigns a lower perplexity to the real proteins, whereas a composition-based method would not be able to distinguish real and synthetic proteins. These results suggests that the RNN-LM perplexity can be used as a threshold to discriminate between real and random proteins.

**Fig. 5:**
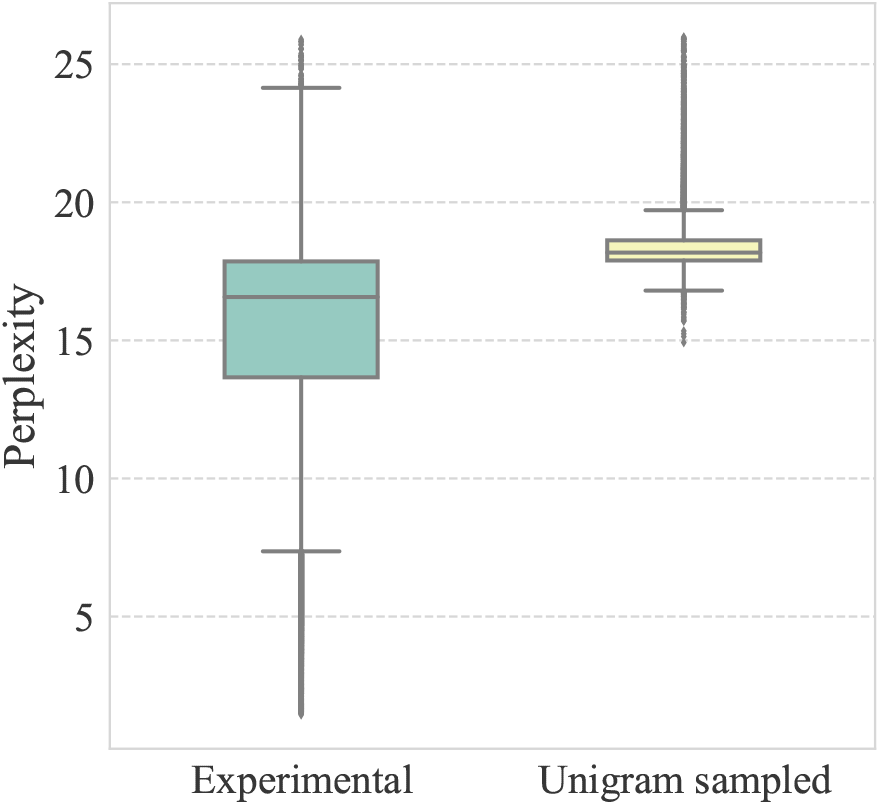
Comparison of the RNN-LM perplexity assigned to experimental eukaryotic proteins and proteins sampled from the unigram distribution.

#### 5.7.3 Generating new proteins

RNN-LM can generate novel proteins, as elaborated in Section 3.4. However, it is challenging to assess the quality of such proteins as we cannot directly confirm whether a generated protein looks like a real protein. The best approach would be to express this protein in a host cell and study its behaviour. As our resources do not extend that far, we analyse the signal peptides (SPs) of proteins generated by the RNN-LM. We observe two interesting facts using this methodology. Firstly, of the 10K generated proteins, 15.62% of them are predicted as having an SP by SignalP 5.0. On the other hand, a set of the same size with random sequences sampled from amino acid frequencies (unigram model) only achieved a 0.66% of predicted SPs. For comparison, the percentage of proteins with SPs in the human proteome is 14.95%. Secondly, we investigate the composition of amino acids in the SPs. In Figure 6 we plot the peptide logos of experimental, LM and random proteins with predicted signal peptides. For random proteins, due to the low amount of signal peptides, the logo plot does not capture the true SP motif whereas the logos for experimental and RNN-LM generated proteins looks indistinguishable.

**Fig. 6:**
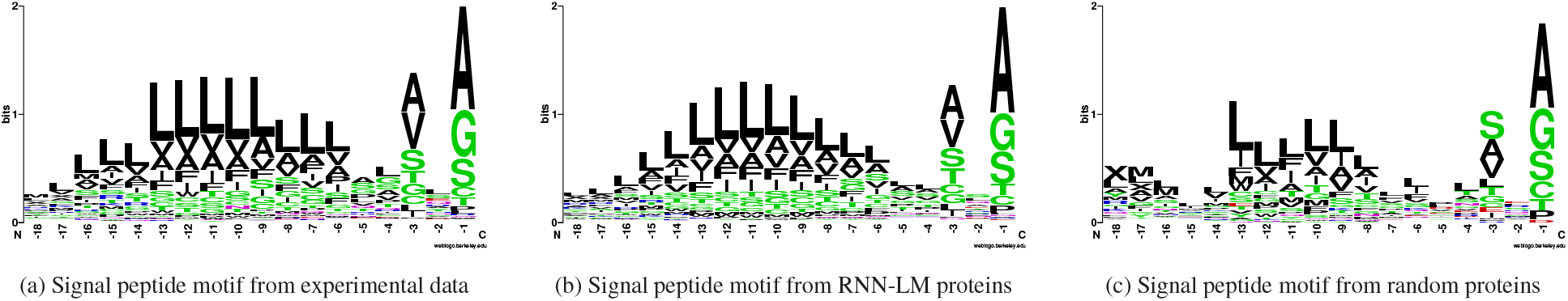
Comparison between signal peptides predicted in experimental test set proteins, novel proteins generated by the RNN-LM from scratch, and proteins that have been randomly generated using the unigram probabilities.

We can thus conclude that the RNN-LM is able to generate realistic secretory proteins by analysing the SP of the generated sequence. However, it still remains unknown whether these proteins can be secreted, folded correctly, and develop any function in or outside the cell.

## 6 Conclusion

In this manuscript, we have studied protein specific dataset properties that can affect the performance of language models (LMs), in particular the Recurrent Neural Network LM (RNN-LM). We have demonstrated that the domain of origin and the experimental evidence of the protein data can have an effect on the perplexity of LMs. The LMs have been tested on a new dataset, with protein specific properties, that we have assembled to be used in future work for benchmarking language models on protein sequences.

By investigating taxonomic domains we find that Bacteria are generally easier to predict while viral proteins are the most difficult to model. Furthermore, incomplete proteins can be as informative as full proteins when training an LM.

In our qualitative analysis, our findings suggest that RNN-LM can generate realistic proteins with a signal peptide distribution that matches what is found in experimentally validated proteins from UniProt.

With this work we aspire to facilitate language modeling research by supplying a stringently formatted dataset. We hope that this can yield better amino acid embeddings for all domains of life and assist improvements in protein prediction tasks.

1 https://github.com/salesforce/awd-lstm-lm

